# Glyphosate exposure minimally alters the *Corbicula fluminea* microbiome while associated bacteria exhibit high tolerance and degradation potential

**DOI:** 10.64898/2026.10.11.758334

**Authors:** Savanna M Eggenberger, Champagne Cunningham, Jessica Perez Godinez, Rachel E Diner

## Abstract

Glyphosate (GLY), the active ingredient in Roundup, is a broad-spectrum herbicide widely used in commercial agriculture and residential landscaping. Due to its extensive application and high water solubility, GLY is frequently introduced into freshwater ecosystems where it has been shown to negatively impact aquatic life and beneficial microorganisms. The freshwater clam *Corbicula fluminea*, a nonnative bivalve species widely distributed throughout the United States, is highly tolerant of many environmental stressors and may serve as a reservoir for microbial taxa capable of tolerating or degrading environmental pollutants. As efficient filter-feeders, these clams are continuously exposed to waterborne contaminants, potentially facilitating interactions between pollutants and host-associated microbial communities.

In this study, we investigated how glyphosate impacts the structure and diversity of *Corbicula fluminea* microbiomes and characterized GLY tolerance and degradation potential in individual microbiome members. We found that microbiome taxonomic composition and abundance-based diversity metrics were minimally impacted by glyphosate, while presence/absence-based phylogenetic diversity shifted depending on treatment concentration.

Furthermore, high GLY concentrations were linked to reduced Faith’s PD alpha diversity, while other alpha diversity metrics were not impacted. Additionally, we isolated and characterized glyphosate-tolerant bacterial strains from clam tissues and assessed their presence within the microbiome. Several isolates exhibited tolerance to very high GLY levels (up to 1600 mg/L), and several also utilized GLY as a sole phosphorus source and possessed the *phnJ* gene, suggesting potential degradation capabilities. While isolated bacteria were minority members of the microbiome, more abundant uncultured taxa were phylogenetically related to known glyphosate degraders. Collectively, these findings suggest that freshwater bivalves may harbor diverse microbial taxa capable of tolerating or degrading herbicides, which may contribute to host resilience in contaminated freshwater environments and potentially represent unexplored avenues for bioremediation research.

## Introduction

Freshwater ecosystems receive contaminants from both agricultural and urban activities (1–3). Herbicides are an important component of this pollution because they can enter waterways through runoff, drainage, and applications near surface waters (4, 5). Glyphosate is among the most widely used herbicides and is frequently detected in freshwater environments influenced by agriculture and development (6, 7). Although designed to target plants, glyphosate and other herbicides can also affect non-target aquatic organisms by altering growth, reproduction, immune function, behavior, and survival (8–10). They may also disrupt microbial activity, nutrient cycling, primary production, and food-web interactions (11–13). Understanding how aquatic organisms respond to glyphosate is therefore important for predicting its broader ecological effects.

Freshwater bivalves are useful models for studying interactions between aquatic organisms and environmental contaminants. Through filter feeding, they process large volumes of water and are repeatedly exposed to dissolved compounds, suspended particles, and associated microorganisms (14). Many bivalves can also accumulate contaminants in their tissues, which has led to their widespread use as sentinel organisms in aquatic monitoring programs (15, 16).

This exposure is ecologically important because freshwater bivalves contribute to water filtration, nutrient cycling, sediment processes, and benthic–pelagic coupling (17, 18). They also serve as prey for wildlife, and some species support commercial, recreational, or subsistence harvests (19, 20). However, many native freshwater bivalves are threatened by pollution and habitat degradation, while some widespread nonnative species persist in highly disturbed environments (21, 22). One such species is *Corbicula fluminea*, an invasive freshwater clam that occupies a broad range of habitats and tolerates substantial environmental variation (23). Its persistence in impacted waterways makes it a useful system for investigating biological traits associated with contaminant resilience.

Despite extensive research on pollutant effects in bivalves, much less is known about how contaminants influence their host-associated microbiomes or how individual microbiome members interact with those compounds. Across animal systems, microbiomes can affect nutrition, immune function, pathogen resistance, and responses to environmental stress (24, 25). Host-associated microorganisms may also modify contaminant exposure by transforming, sequestering, or degrading pollutants (26, 27). These interactions may be especially relevant for glyphosate because its molecular target, 5-enolpyruvylshikimate-3-phosphate synthase, is present in plants and many microorganisms but absent from animals (28, 29). Glyphosate may therefore influence bivalves indirectly through effects on associated microbial communities. At the same time, some bacteria can tolerate high glyphosate concentrations or metabolize the compound through pathways involving glyphosate oxidoreductase or carbon–phosphorus lyase (30, 31). Previous studies have examined pollutant-associated changes in microbiome structure, and others have characterized bacterial isolates capable of contaminant tolerance or degradation (32–34). However, few studies have integrated these approaches to determine whether community-level responses are linked to the functional traits of individual microbiome members. This gap limits our understanding of how microbial composition and functions jointly shape host responses to environmental pollution.

In this study, we used the invasive freshwater clam *Corbicula fluminea* as a model for investigating microbiome responses to glyphosate exposure. *Corbicula fluminea* is broadly distributed, inhabits waterways influenced by agricultural and urban activity, and tolerates a wide range of environmental conditions (23, 35). We therefore hypothesized that its microbiome would be relatively resilient to glyphosate and would contain bacterial taxa capable of tolerating or potentially degrading the herbicide. To investigate this, we conducted a controlled glyphosate exposure experiment to examine microbiome changes using16S rRNA gene amplicon sequencing, and subsequently isolated and characterized clam-associated bacteria and conducted glyphosate tolerance assays, phosphorus-utilization assays, and screening for the *phnJ* gene, which is often involved in microbial glyphosate degradation. Specifically, we asked whether glyphosate exposure alters clam microbiome composition and diversity; whether individual clam-associated bacteria are sensitive or tolerant to glyphosate and how prevalent tolerant isolates are within the microbiome; and which cultured or uncultured microbial taxa may be capable of glyphosate degradation. By linking community-level patterns with isolate-level phenotypes, this study provides insight into microbial traits that may contribute to bivalve persistence in contaminated environments. It also evaluates freshwater bivalve microbiomes as a potential source of microorganisms relevant to pollutant degradation and bioremediation.

## Materials and Methods

### Corbicula fluminea glyphosate exposure experiment

*C. fluminea* clams were collected from the Arthur Halle Stadium site at Nonconnah Creek in Memphis, Tennessee using hand tools and transported on ice back to the University of Memphis. The clams were then scrubbed with sterile brushes, rinsed with reverse osmosis (RO) water, and sprayed with 75% ethanol to reduce environmental contamination. Clams were acclimated to laboratory conditions in a 10-gallon tank with filtered (0.2 µm), aerated tap water (adjusted to pH 7) for 7 days prior to the exposure experiment. Water was changed every 48 hrs. Clams were fed daily with a green algae mixture of *Chlorella vulgaris* and *Chlamydomonas reinhardtii* at a 1:3 ratio with a final concentration of 800,000 cells/clam/day. After acclimation, eighteen clams were split into 3 different treatments: control (no glyphosate), low glyphosate (100 µg/L), and high glyphosate (2 mg/L). Each treatment consisted of 3 aerated 2 L jars with 2 clams in each, resulting in N=6 clams per treatment. Water was changed daily with replenished glyphosate concentrations, and clams continued to be fed daily. After 4 days of exposure clams were harvested and shucked with a sterile scalpel. Whole clams (∼0.3-0.5 g of wet tissue) were manually homogenized with a tissue grinder in 1.5 mL tubes with 600 µl Zymo DNA/RNA shield solution and stored at –20°C prior to microbiome analysis.

### Isolation and culturing of clam-associated bacteria

*C. fluminea* clams were collected and cleaned as described above. Five clams were opened using scalpels and dissected to collect foot, gill, and digestive gland tissues. Tissue samples (∼25-50 mg of tissue) were preserved in 600 µl Zymo RNA/DNA Shield solution at - 20°C for downstream microbiome analysis. For bacterial isolation, 25-50 mg of each tissue type was added to a 1.5 mL microcentrifuge tube containing 0.5 mL of sterile zirconia/silica beads (1 mm diameter, biospec) and 500 µl sterile phosphate buffered saline (PBS). Samples were vortexed on a Vortex Genie machine for 5 mins on the highest setting (10). 100 µL of supernatant was then removed and serially diluted (1:10, 1:100, 1:1000, 1:10000) using sterile PBS. 100 µl from each dilution was spread on Luria-Bertani (LB) agar plates and incubated at 30°C for ∼24 hrs, after which individual colonies were streaked on fresh LB plates and incubated under the same conditions to confirm single colonies. Resulting isolates were grown overnight in LB broth before being preserved in a glycerol stock solution (25 % final concentration glycerol) and stored at –80°C. 105 isolates were successfully re-grown after freezing and used in subsequent glyphosate exposure screens.

To assess isolate sensitivity to glyphosate, LB agar plates were enriched using a 0.2 µm filter-sterilized 1 g/L glyphosate stock solution (Thermo Scientific) to achieve final concentrations of 100 mg/L, 200 mg/L, 400 mg/L, 800 mg/L, and 1600 mg/L. Bacterial isolates were revived from glycerol stocks in LB and incubated overnight under shaking conditions (250 rpm, 30°C). Optical densities at 600 nm (OD600) were measured the following day using spectrophotometry and adjusted to a uniform OD600 of 0.1 before being further diluted 1:100 using PBS buffer. A total of 10 µl from the diluted cultures was then spot plated on control (LB only) and LB/GLY-enriched plates in a grid formation and incubated at 30°C in the dark for 48 hrs. Growth was then assessed and isolates displaying growth at the 1600 mg/L concentration (10 total) were selected for further testing.

To assess the ability of the ten highly tolerant isolates to utilize glyphosate as a sole phosphorous source, a modified MOPS based minimal media (36) was prepared containing (g/L): 10.47 MOPS, 2.47 (NH4)2SO4, 0.5 K2SO4, 0.08 MgSO4 anhydrous, 0.08 CaCl2 * 2H2O, in addition to 10 mL trace metals solution, 25 mL glucose, 80 mL glyphosate stock solution (5 g/L) and molecular grade H2O. The pH of the MOPS/GLY medium was adjusted to 7.2 and filter-sterilized (0.2 µm) prior to use. Inoculant was collected from the LB/GLY 1600 mg/L plates, transferred to 3 mL LB cultures, and grown overnight at 30°C, 250 rpm. Cells were washed 3 times using MOPS media and centrifuged at 12,000 x g for 2 mins. The supernatant was then removed, and the cells were resuspended in new MOPS media. OD600 of the washed cell suspensions were measured and adjusted to 0.01 nm. Cultures were grown in triplicate glass tubes along with an *E. coli* DH5α strain serving as a negative control due to its known inability to degrade glyphosate (37). Samples were incubated in the dark under shaking conditions (250 rpm, 30°C), and bacterial growth was measured at OD600 at 24-hr intervals over a 7-day period. 1 mL of each culture was collected for a second passage, and cells were washed and standardized as described above before being inoculated into fresh media.

### Sanger sequencing and analysis of 16S rRNA and phnJ genes

To identify the bacterial isolates most tolerant to glyphosate, the 16S rRNA gene was amplified using DreamTaq 2X PCR Master Mix (Thermo Scientific) in 25 µl reactions following the manufacturer recommended protocol (Ta= 58°C) with the universal primer pair 27F (5′-AGAGTTTGATCCTGGCTCAG-3’) and 1492R (5′-TACGGTTACCTTGTTACGACTT-3′).

Successful amplification was confirmed via gel electrophoresis, purified using the Zymo Clean and Concentrate –25 kit, and sequenced by Sanger Sequencing with Eurofins Scientific. A consistent 1000 bp region beginning at nucleotide 20 was selected for analysis (Full sequences available in Supplemental Table 1). The *phnJ* gene, which is frequently involved in glyphosate degradation (37, 38) and has an expected amplicon size of 550-650 bp was also amplified using DreamTaq 2X PCR Master Mix and the primer set Fphnj202 (5’-AGACCCGYCACCGCATCC-3’), Rphnj201 (5’-CAATARTCGGTRTCGGAGCAGAC-3’)(39). The PCR was performed in 30 cycles as follows: initial denaturation step at 95°C for 2 mins, second denaturation step at 95°C for 45 sec, annealing step at 60°C for 45 sec, extension step at 72°C for 30 sec, and final extension step at 72°C. PCR confirmation and sequencing was carried out as above. A consistent 450 bp region beginning at nucleotide 20 was used for analysis (Full sequences available in Supplemental Table 1).

The Basic Local Alignment Search Tool (BLAST) developed by the National Center for Biotechnology Information (NCBI) was used to annotate sequences and provide percent identities for bacterial taxonomic assignments. Additionally, *phnJ* sequences were aligned using CLUSTALW and an unrooted phylogenetic tree was constructed using the sequence of an *E. coli* K12 strain in addition to (5) other *phnJ* sequences found in bacteria with confirmed glyphosate degradation abilities.

### Microbiome sequencing

For the glyphosate exposure experiment, ∼25-50 mg of homogenized whole clam tissue from the storage shield solution was used for DNA extraction with the Zymo DNA Miniprep Kit. Following quantification via nanodrop, DNA was stored at –20°C until sequencing. Foot, gill, and digestive gland samples (∼25-50 mg) collected from the five clams that were also used for bacterial isolations and subsequent glyphosate sensitivity assays were preserved in 600 µl of Zymo DNA/RNA shield at-20°C. Tissues were later homogenized with a tissue grinder and DNA extractions performed using the ZymoBIOMICS DNA Miniprep Kit. Purified DNA samples were sent for 16S rRNA gene amplicon sequencing of the V3-V4 region at either the University of Mississippi Molecular and Genomics Core (glyphosate exposure experiment) or Zymo (tissue-specific microbiomes). Sequences were returned as demultiplexed paired end reads for downstream bioinformatics analysis.

### Microbiome Bioinformatic and Statistical Analyses

For both the tissue-specific and glyphosate exposure microbiome sequences, the QIIME2 pipeline (version qiime2-amplicon-2025.10)(40) was utilized for quality control, trimming, and diversity and taxonomic analyses. DADA2 (41) was used to trim primers, merge sequences, and assign ASVs. Taxonomic assignment of ASVs was performed using a naive Bayes classifier trained on the Silva version 138 database (42) and mitochondrial, chloroplast, eukaryotic, and unassigned domain sequences were removed prior to further analysis. Taxonomy bar plots were generated for both microbiome data sets to visualize microbiome community composition. Alpha diversity analyses were performed with Faith’s Phylogenetic Diversity (Faith PD), Shannon diversity, Evenness, and observed ASV metrics after rarefaction to a standardized sequencing depth (tissue-specific = 43,000; glyphosate exposure = 300,000), and outputs were subsequently visualized using R (43). For beta diversity analyses Bray-Curtis, Jaccard, unweighted UniFrac, and weighted UniFrac metrics were examined using QIIME2 core-metrics-phylogenetic with the aforementioned sampling depths, while robust Aitchison Principal Component Analyses (RPCA) were conducted using Gemelli with no rarefaction (44). Statistical significance between groups was determined using pairwise multivariate analysis of variance (PERMANOVA), and principal coordinates analysis (PCoA) plots were generated to visualize microbial community differences across groups. Permutational Analysis of Multivariate Dispersions (PERMDISP) was conducted in QIIME2 to assess dispersion of samples within groups, which determined dispersal of all groups to be homogeneous (q > 0.05 for all pairwise comparison). Analysis of Composition of Microbiomes with Bias Correction (ANCOM-BC2)(45) was used to conduct differential abundance analyses for microbiome datasets and applies multiple-testing correction (significance = q < 0.05) and sensitivity analysis.

Negative and positive control samples were sequenced alongside microbiome samples. For the tissue microbiome analysis, quantitative PCR of the 16S rRNA gene was conducted for each sample and used to inform decontamination procedures. Positive controls were the ZymoBIOMICS Microbial Community DNA Standard and, following microbiome analysis, taxonomy bar plots were examined to confirm the expected microbiome abundance composition described by the manufacturer. For negative controls, Zymo RNA/DNA shield solution was processed in the same manner as biological samples with the Zymo DNA miniprep kit. To assess and account for potential contamination, blank samples and controls were visually and statistically compared using the filtered tables with mitochondrial, chloroplast, eukaryotic, and unassigned domain sequences removed (noECMU). In beta-diversity PERMANOVAs and PCoA plots, taxonomy bar plots, and number of observed ASVs, blank communities were distinct from samples in both datasets. Next, we identified potential contaminants using QIIME quality-control decontam-identify. For the glyphosate exposure experiment, we employed the prevalence method and did not identify any contaminant ASVs requiring removal, so the original noECMU tables were used for the final analysis. For the tissue-specific microbiome, a combined prevalence/frequency approach was used, and features were flagged if they were enriched in negative controls and/or showed the expected inverse relationship with 16S copy concentration.

Candidate contaminants were filtered from the feature table using a threshold of p < 0.1, then downstream analyses were rerun on the decontaminated table. The full suite of microbiome analyses were run on both the pre-filtered and post-filtered datasets revealing no significant differences in any metric.

## Results

### Effect of Glyphosate on the Corbicula fluminea Microbiome

Experimental treatment was associated with significant differences in Faith’s phylogenetic diversity (alpha diversity) both overall (Kruskal-Wallis p = 0.0043), and in pairwise Wilcoxon tests followed by Benjamini-Hochberg correction which showed differences in no GLY vs. low GLY (q = 0.045), no GLY vs. high GLY (q = 0.025), and low GLY vs. high GLY (q = 0.045)(Supplemental Figure 1). In contrast, other alpha diversity metrics including Shannon diversity, evenness, and observed ASVs did not differ significantly among treatments. Beta diversity results depended on the distance metric used. Bray-Curtis, Jaccard, weighted UniFrac, and robust Aitchison RPCA analyses (Figure 1A) did not show significant treatment effects.

**Figure 1.**
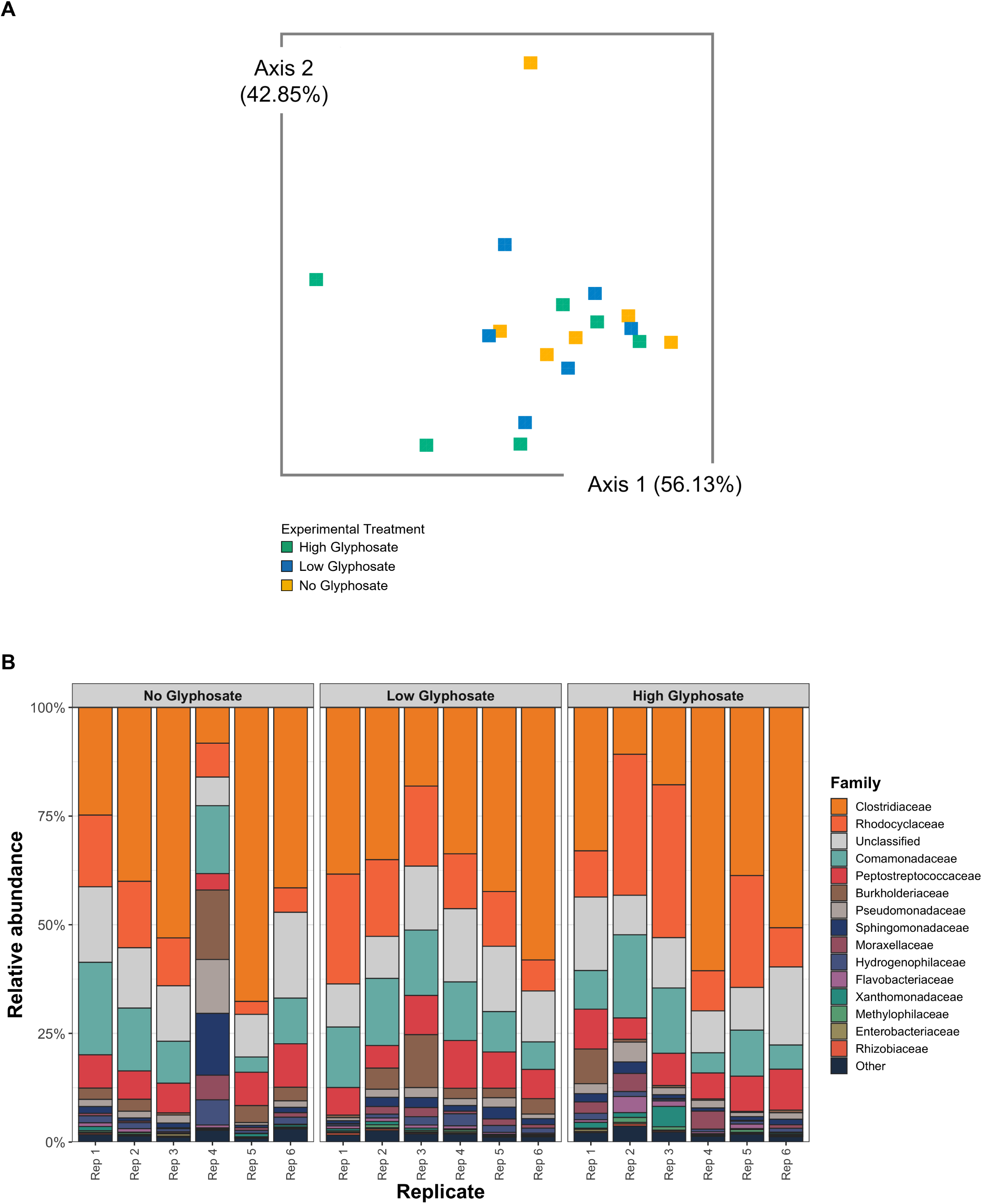
A) Emperor biplot visualizing microbiome beta diversity from glyphosate exposure experiment using robust Aitcheson RPCA analysis. Whole clam tissues were used for microbiome analysis. B) Family-level taxonomy bar plot for *Corbicula fluminea* microbiomes assessed during glyphosate exposure experiment. Each bar represents an individual biological replicate. Samples are from homogenized clams.

Taxonomy bar plots showed a similar profile apart from replicate 4 within the no GLY treatment (Figure 1B). However, unweighted UniFrac distances differed significantly among treatment groups, specifically pairwise unweighted UniFrac comparisons showed significant differences between high and low GLY treatments (q = 0.0075) and between high and no GLY treatments (q = 0.0075), while low GLY and no GLY did not differ significantly. Results of all PERMDISP analyses were not significant (p <0.05). ANCOM-BC2 identified one differentially abundant taxon at the genus level: *Sphaerotilus* was significantly depleted in the high GLY treatment relative to the no GLY control (log fold change =-4.50, q = 0.014). No significant differentially abundant ASVs or families were detected.

### Tissue-Specific Microbiome Profiling

Microbial diversity differed only weakly among tissue types. Alpha diversity metrics were generally similar among foot, gill, and digestive gland samples, with no significant differences detected for Faith’s phylogenetic diversity, Shannon diversity, Evenness, or observed ASVs either across samples or in pairwise comparisons (Supplementary Figure 2). Beta diversity analyses included in the core diversity metrics did not show overall separation of tissue-associated bacterial communities: PERMANOVA tests based on Bray-Curtis (p = 0.346), Jaccard (p = 0.791), unweighted UniFrac (p = 0.835), and weighted UniFrac (p = 0.293) distances were not significant, and PERMDISP tests were also non-significant, indicating no evidence that group dispersion explained the observed patterns. Robust Aitchison RPCA likewise was not significant in the overall PERMANOVA (p = 0.230), although the pairwise RPCA comparison between foot and digestive gland samples was significant after correction (q = 0.015) (Supplemental Figure 3A). Additionally, taxonomy bar plots did not show clear differentiation between tissue types (Supplemental Figure 3B). ANCOM-BC2 did not identify any differentially abundant taxa among tissue types at the ASV, genus, or family level after multiple-testing correction and sensitivity analysis.

### Isolation and Screening of Glyphosate Tolerant Bacteria

A total of 105 individual isolates were obtained from serial dilutions of clam foot, gill, and digestive gland tissues and screened for glyphosate sensitivity at concentrations of 100, 200, 400, 800, and 1600 mg/L. All 105 isolates exhibited growth at 100 mg/L glyphosate. Growth remained common at 200 and 400 mg/L, with 104 (99.05%) and 100 (95.24%) isolates growing, respectively. At 800 mg/L, growth declined substantially, resulting in 82 successful isolates (78.10%). Only ten isolates overall (9.52%) were capable of growth at 1600 mg/L. Of these highly glyphosate-tolerant isolates, eight originated from foot tissues, while one each originated from digestive gland and gill tissues. These ten isolates were selected for further testing.

### Isolate Growth with Glyphosate as Sole Phosphorus Source

Based upon the growth trends within the first and second passages, 8 isolates showed significant growth in MOPS/GLY 400 mg/L (Figure 2), indicating efficient use of glyphosate as a sole phosphorus source. Iso-89 and Iso-99 did not grow considerably within either passage, with OD600 values remaining similar to those of the *E. coli* control (< 0.03 nm). All bacteria with growth ability required a lag period during passage 1, which was highly variable between isolates and ranged between ∼24-120 hrs. Iso-88 exhibited the highest OD600 values in passage 1, reaching values > 0.29 nm despite having a lag period of ∼96 hrs. Similarly, Iso-42 and Iso-111 did not reach an exponential growth period until ∼120 hrs but reached > 0.15 nm. Most isolates did not require a sustained lag period during passage 2, with five reaching exponential growth within the first 24 hrs and reaching higher optical densities than achieved during passage 1. Iso-42 and Iso-111 showed the highest growth density during passage 2, both reaching > 0.30 nm. Iso-100, which did not grow significantly during passage 1, did reach exponential growth between 48-72 hrs.

**Figure 2.**
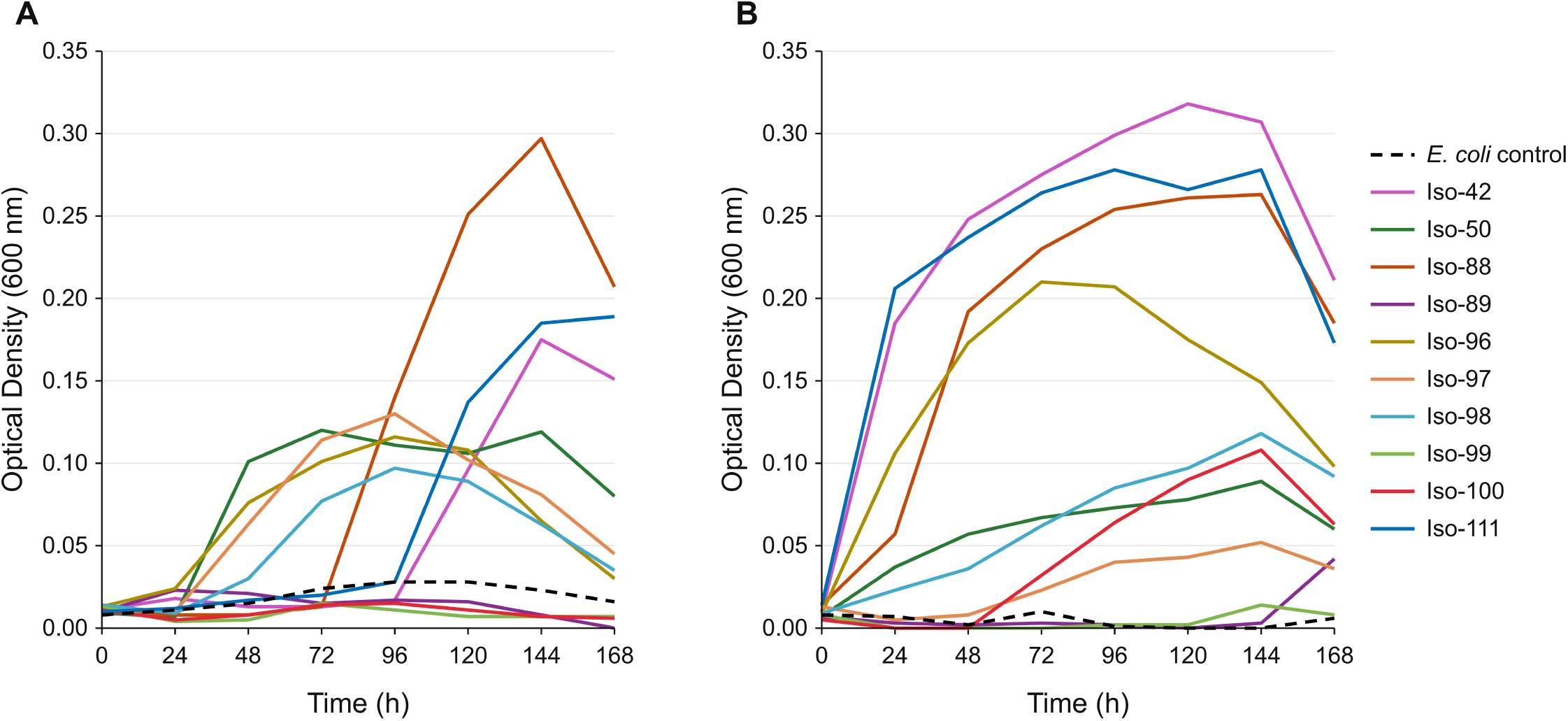
Growth of bacterial isolates cultured from *Corbicula fluminea* in MOPS media containing 400 mg/L glyphosate as the sole phosphorus source. Growth was measured using optical density at 600 nm (OD600) over time. A) Passage 1 B) Passage 2. Cultures were standardized and transferred to fresh MOPS media supplemented with glyphosate 400 mg/L between passages.

### Bacterial Identification and phnJ Gene Detection

Analysis of the 16S rRNA gene sequences revealed that all ten isolates belonged to one of two major bacterial families: *Enterobacteriaceae* and *Yersiniaceae* (Table 1). Iso-98, Iso-99, and Iso-100 were most closely related to *Rahnella aquatilis,* while Iso-42 and Iso-111 showed the highest similarity to *Lelliottia amnigena*. Although the *R. aquatilis* and *L. amnigena* isolates showed a high level of similarity to each other, there were nucleotide variations within the sequences indicating distinct strains. The remaining isolates were most closely related to *Ewingella americana* (Iso-50), *Citrobacter freundii* (Iso-88)*, Atlantibacter hermannii* (Iso-89), and *Klebsiella pneumoniae* (Iso-96). Iso-97 could not be confidently identified (87.69% identity) but demonstrated closest similarity to *Pantoea agglomerans*. Phylogenetic analysis of 16S rRNA gene sequences showed that isolates were closely related overall, with isolates of the same species clustering together (Figure 3A). Additionally, reference sequences from known glyphosate degrading taxa clustered together.

**Figure 3.**
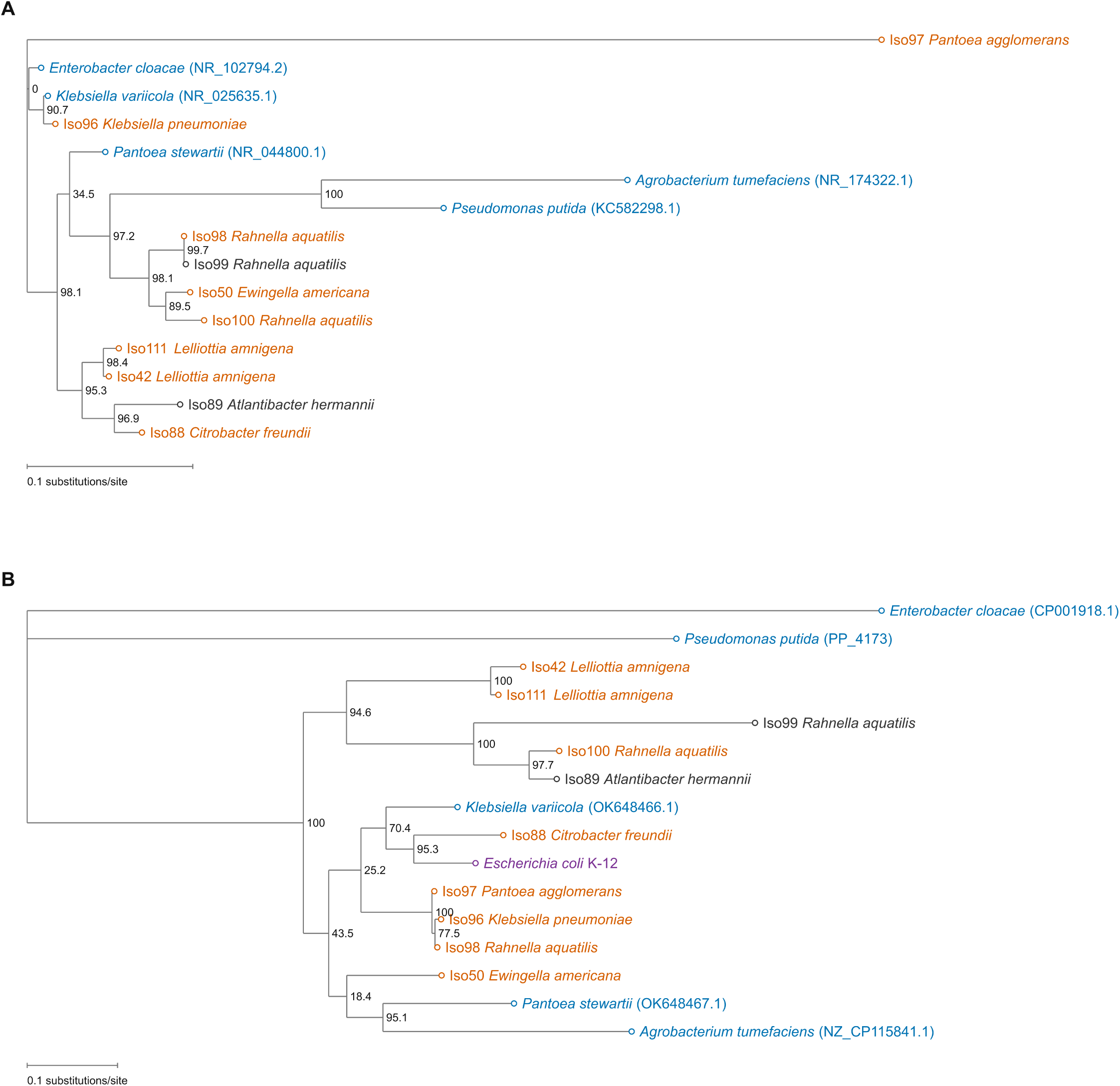
Unrooted phylogenetic trees based on (A) 16S rRNA and (B) phnJ gene sequences. Node values indicate SH-like local support on a 0-100 scale, with higher values indicating stronger support. Blue labels indicate known glyphosate-degrading references; orange indicates isolates that grew on glyphosate; gray indicates isolates that did not grow on glyphosate; and purple identifies the *E. coli* reference. Scale bars represent 0.1 substitutions per site.

**Table 1.** Identification and glyphosate-degradation associated characteristics of bacterial isolates cultured from *Corbicula fluminea* foot, gill, and digestive gland tissue. Isolates were identified using 16S rRNA gene Sanger sequencing, with percent identity representing sequence similarity to closest NCBI match. Growth on glyphosate indicates the ability to use glyphosate as a sole phosphorus source. Glyphosate degradation indicates whether identified species have been associated with glyphosate degradation in literature; supporting references are provided where applicable.

| Isolate ID | Closest species-level identification | 16S rRNA identity (%) | Growth on Glyphosate | Prior evidence of GLY degradation | Reference(s) |
| --- | --- | --- | --- | --- | --- |
| Iso-42 | *Lelliottia amnigena* | 99.6 | Yes | No | - |
| Iso-50 | *Ewingella americana* | 98.7 | Yes | No | - |
| Iso-88 | *Citrobacter freundii* | 98.88 | Yes | Yes | Hamieh et al. 2025; Owuna et al. 2021 |
| Iso-89 | *Atlantibacter hermannii* | 98.46 | No | No | - |
| Iso-96 | *Klebsiella pneumoniae* | 99.6 | Yes | Yes | Hamieh et al. 2025; Knecht et al. 2026 |
| Iso-97 | *Pantoea agglomerans* | 87.69 | Yes | Yes | Maldani et al. 2018; Masotti et al. 2021 |
| Iso-98 | *Rahnella aquatilis* | 99.9 | Yes | Yes | Benslama and Boulahrouf 2013; Peng et al. 2012 |
| Iso-99 | *Rahnella aquatilis* | 99.8 | No | Yes | Benslama and Boulahrouf 2013; Peng et al. 2012 |
| Iso-100 | *Rahnella aquatilis* | 99.6 | Yes | Yes | Benslama and Boulahrouf 2013; Peng et al. 2012 |
| Iso-111 | *Lelliottia amnigena* | 99.5 | Yes | No | - |

*Phnj* gene sequences were successfully amplified for all 10 isolates and matched to previously annotated C-P lyase *phnj* genes in the NCBI database. This supports the presence of *phnj*-like genes within these isolates, though presence of the gene does not necessarily support gene function. CLUSTALW alignment revealed that isolates contained divergent *phnJ* sequences, which was also supported by the phylogenetic tree (Figure 3B). Notably, *R. aquatilis phnJ* sequences for Iso-98 (growth on glyphosate) and Iso-99 (no growth on glyphosate) showed substantial divergence from each other despite high 16S rRNA sequence identity and close phylogenetic placement. Sequences showed variable relationships to reference sequences of known glyphosate degrading taxa, with several isolates clustering near references.

### Assessing the prevalence of potential glyphosate degraders in C. fluminea microbiomes

Combining the microbiome datasets and bacterial isolate glyphosate assays, we examined the prevalence of glyphosate-tolerant and potential glyphosate degrading taxa within the *C. fluminea* microbiome. By performing a literature search to compile a list of known glyphosate degrading taxa and comparing this to our microbiome abundance data, we identified 14 groups of bacteria at the genus level that are known to degrade glyphosate or utilize it as a nutrient source. These 14 included *Acidovorax*, *Bacillus*, *Bradyrhizobium*, *Chryseobacterium*, *Citrobacter*, *Comamonas*, *Enterobacter*, *Geobacillus*, *Klebsiella*, *Lysinibacillus*, *Novosphingobium*, *Pantoea*, *Pseudomonas,* and *Rahnella*.

We next calculated the percent relative abundance of the ten previously studied bacterial isolates within the tissue-specific microbiome they originated from (Supplemental Table 2). By comparing the isolates’ 1000 bp sequences to the microbiome ASV sequences, we were able to determine that these isolates were relatively rare members of the *C. fluminea* microbiome (< 1% relative abundance). We identified exact sequence matches for Iso-42 (*L. amnigena*), Iso-98 (*R. aquatilis*), Iso-99 (*R. aquatilis*), and Iso-111 (*L. amnigena*). For the additional five isolates (not including Iso-97 with low sequence quality), we were able to obtain their closest ASV match (>97 % match quality). For the exposure microbiome data set, proxy ASVs with high similarity (>97 %) to the isolate sequences were identified and compared among exposure treatment groups. As with the tissue microbiome comparison, these isolates represented a considerably rare (<1 %) subset of the microbiome. While relative abundances of these taxa were low and did not differ significantly between glyphosate exposure treatments based on ANCOMBC-2 analysis, qualitative observations indicate that some of these taxa had highest relative abundance in the high GLY treatment.

## Discussion

### Limited effects of glyphosate and tissue type on C. fluminea microbiome community diversity and composition

Glyphosate exposure produced detectable but relatively limited changes in the *Corbicula fluminea* microbiome. Faith’s phylogenetic diversity and unweighted UniFrac distances differed among treatments, whereas Shannon diversity, evenness, observed ASVs, Bray-Curtis, Jaccard, weighted UniFrac, and robust Aitchison analyses did not. Together, these results suggest that glyphosate primarily affected the phylogenetic composition of less abundant community members rather than causing broad shifts in dominant taxa or overall abundance structure. The depletion of only one genus, *Sphaerotilus*, in the high-glyphosate treatment further supports the relative stability of the community. Similar selective or metric-dependent microbiome responses to glyphosate have been reported in other aquatic hosts and environmental systems, although the magnitude and direction of these effects vary with dose, exposure duration, host species, and environmental context. For instance, glyphosate has been found to significantly decrease alpha diversity in the rainbow trout *Oncorhynchus mykiss* gill microbiome during periods of chronic exposure (46); however, in the mediterranean mussel *Mytilus galloprovincialis*, glyphosate exposure caused community restructuring at the individual taxa level but did not significantly alter overall bacterial richness (47). The limited response observed here may reflect active host regulation of the microbiome, functional redundancy among microbial taxa, or prior adaptation of *C. fluminea* and its associated microorganisms to chemically disturbed habitats. Bivalves can maintain relatively stable microbial communities despite environmental fluctuations, potentially because these communities contribute to nutrition, immune function, and stress tolerance (48, 49). Such stability may provide adaptive potential under pollutant exposure (50), although the present data does not establish whether microbiome resilience directly improves host performance.

Microbiome composition also differed only weakly among the foot, gill, and digestive gland. Most alpha-and beta-diversity metrics showed no significant tissue effects, and no taxa were differentially abundant among tissues. The only significant comparison was the robust Aitchison RPCA contrast between foot and digestive gland samples. This limited tissue differentiation was somewhat unexpected because bivalve organs differ in physiology, environmental exposure, and function, and stronger separation between gill and digestive tissues has been reported in oysters and other bivalves. For example, a study comparing the gill, gonad, byssus gland, hemolymph, mantle adductor muscle, mucus, and gut microbiomes of the black-lipped pearl oyster *Pinctada margaritifera* found that richness of Operational Taxonomic Units (OTUs) and phylogenetic diversity differed significantly between tissues (51). This differentiation has also been observed in Eastern oysters (52), Pacific oysters (53–55), and Manila clams (56). Several factors may have reduced tissue-specific patterns in this study. These include continual microbial exchange among organs, shared exposure to filtered water and food particles, limited sample size, or substantial variation among individual clams. Microbiome composition in bivalves may also be shaped by season, diet, reproductive state, parasite burden, chemical exposure and local environmental conditions (54, 57, 58). The apparent interindividual variability observed across tissues therefore warrants further investigation. Longitudinal sampling and larger datasets that incorporate host condition, food availability, seasonal change, and parasite loads could help determine whether tissue-specific communities are consistently maintained or are instead highly dynamic within *C. fluminea*.

### Glyphosate-tolerant bacteria associated with C. fluminea demonstrate potential for glyphosate utilization via the C-P lyase pathway

Glyphosate tolerance was high among the bacterial isolates obtained from *C. fluminea* and eight isolates exhibited the ability to utilize it as a phosphorus source. Many of the glyphosate-tolerant isolates identified in this study have previously been associated with environments experiencing frequent exposure or directly linked to glyphosate degradation, suggesting that recovery of these taxa from *C. fluminea* exposed to glyphosate within the aquatic environment is not unexpected. For example, *C. freundii* originating from agricultural soils has been isolated in multiple studies where glyphosate enrichment was utilized for isolation (59, 60). *Klebsiella* and *Pantoea* are both genera that are frequently isolated from glyphosate exposed environments, both terrestrial and freshwater (39). *K. pneumoniae* has also demonstrated glyphosate degradation potential when isolated from glyphosate-applied soils (61) and *P. agglomerans* has been shown to degrade glyphosate-based herbicides (62). Likewise, multiple studies have shown the ability of *R. aquatilis* to utilize glyphosate as a phosphorus source (63, 64). In contrast, *L. amnigena*, *E. americana*, and *A. hermannii*, according to current knowledge, represent novel species with the ability to tolerate glyphosate. The species that exhibited the highest OD600 measurements within this study, *L. amnigena*, has no history of herbicide degradation; however, recent studies demonstrate its ability to perform nitrate remediation (65).

*E. americana,* despite no known evidence of glyphosate degradation, grew utilizing glyphosate in the current study and has previously been shown to efficiently degrade phenol for use as a carbon source (66). Similarly, *A. hermannii*, which showed reduced sensitivity to glyphosate but was unable to use it as a phosphorus source, has been linked to degradation of other compounds, such as crude oil (67) and chlorpyrifos (68). Therefore, our results support the findings of previous studies but also identify additional species capable of tolerating glyphosate, some of which demonstrated glyphosate utilization and may provide candidate taxa for future glyphosate degradation studies.

The *phnJ* gene, responsible for the cleavage of the carbon-phosphorus bond of phosphonates (38), was detected in all ten isolates. The detection of *phnJ* in these isolates provides further evidence for their glyphosate-utilization potential via the C-P lyase pathway. The increased occurrence of phosphonate-utilization genes among environmentally associated bacteria is thought to reflect the needs of these bacteria to overcome phosphorus limitations (69). Thus, the presence of phosphonate-utilization genes among the isolates within this study may reflect their capacity to access alternative phosphorus sources under limited conditions. Previous studies have demonstrated C-P lyase-mediated phosphonate utilization in *K. pneumoniae* (70), and *phnJ* is also present in publicly available genome annotations for *K. pneumoniae*, *C. freundii* (71), and *L. amnigena* (65). The presence of *phnJ* alone cannot definitively establish a functional C-P lyase pathway or prove glyphosate degradation, a distinction that was apparent among our isolates, as the gene was detected in all ten while only eight demonstrated growth with glyphosate as the phosphorus source. Furthermore, *E. coli*, which was used as a negative control in this study, possesses the *phnJ* gene but is not capable of glyphosate degradation due to inefficient phosphonate transport (37). In one of the *R. aquatilis* strains (Iso-99), *phnJ* was present but did not correspond to growth on glyphosate, while two other bacterial taxa identified as the same species (Iso-98, Iso-100) successfully grew. The *phnJ* sequence of Iso-99 was highly divergent from the other two strains, indicating the possibility of strain-level variation in C-P lyase functionality which is supported by previous studies (72). These differences could be associated with loss of functionality in the *phnJ* gene itself, differences within the *phn* operon, regulatory differences that prevent expression, or deficient phosphonate uptake similar to *E. coli*.

The same explanations could account for the inability of *A. hermannii* to grow under these conditions, despite also possessing the *phnJ* gene. Our results provide genetic evidence supporting phosphonate-utilization potential among these bacteria, while growth assays demonstrated the functional capacity for glyphosate utilization as a phosphorus source, supporting involvement of the C-P lyase pathway. However, additional studies incorporating glyphosate reduction rates alongside bacterial growth measurements are required to confirm glyphosate degradation.

### Glyphosate degradation potential extends beyond cultured isolates within the C. fluminea microbiome

While the ten isolates identified in this study represent minority members of the *C. fluminea* tissue microbiomes, other more abundant taxa belonging to groups with previously documented glyphosate degradation capabilities were identified within the exposure and tissue-specific microbiome datasets. Additionally, these more abundant taxa may have been represented in the 105 isolates that demonstrated high GLY tolerance during the sensitivity screening, just not surviving at the highest concentration. Thus, these cultured isolates likely represent only a small fraction of bacteria within *C. fluminea* with potential glyphosate-utilization or degradation capabilities. For example, the *Pseudomonas* genus, which represented several taxa within the microbiome, contains multiple well-characterized glyphosate degrading species, including *Pseudomonas putida* (63) and *Pseudomonas aeruginosa* (73). Degradation has also been demonstrated in members of other taxa present within the *C. fluminea* microbiome, including *Novosphingobium* (74), *Comamonas* (75), and *Acidovorax* (74). The discrepancies between taxa detected through amplicon sequencing and those recovered from tissues were to be expected.

Culture-dependent methods are unable to recover all host-associated bacteria, as many require specific nutrients and environmental conditions to survive (76). Our initial isolation methods using LB agar may have favored rapidly growing, metabolically versatile taxa, contributing to the predominance of *Enterobacteriaceae* and *Yersiniaceae* among our isolates (77, 78); therefore, the absence of these taxa among our isolates should not be interpreted as evidence that they lack a role in glyphosate processing within the *C. fluminea* microbiome. Future studies should incorporate glyphosate-enrichment and additional media types during isolation to increase the probability of recovering majority members of the clam microbiome.

Our findings of glyphosate-utilizing bacteria from *C. fluminea*, along with the presence of taxa associated with glyphosate degradation within the microbiome, suggest that bivalve-associated microbial communities may represent another reservoir for bacteria with these metabolic capabilities. The presence of bacteria with potential glyphosate degradation capabilities within the *C. fluminea* microbiome is consistent with previous evidence in other systems. For example, highly glyphosate-tolerant bacteria have been isolated from poultry (79) and from farmers frequently exposed to the herbicide through their work (80). Occurrence across host systems suggests that the observations of this study are not unique to *C. fluminea* alone, but instead reflect a broad distribution of bacteria capable of tolerating the herbicide. However, the efficient filter-feeding capabilities of *C. fluminea* and other bivalve species, and their ability to concentrate herbicides within tissue, may lead to more efficient removal of contaminants from the freshwater ecosystems they inhabit compared to others. Glyphosate-degrading bacteria are similarly found in aquatic environments, including lakes, rivers, creeks, and sediments (39) and can significantly reduce contamination (81). These findings are particularly relevant because microbial degradation represents a vital route through which glyphosate can be broken down and transformed following its entry into water bodies. Although this study was limited in that glyphosate degradation rates/removal were not quantified, these results support the need for further investigation into the use of bivalves and their associated bacteria as bioremediation candidates. Further studies incorporating direct measurements of glyphosate degradation could elucidate whether individual isolates or mixed microbial consortia from bivalves can effectively remove glyphosate from contaminated aquatic environments.

## Conclusion

This study provides an integrated assessment of the *C. fluminea* microbiome and its potential for harboring glyphosate-tolerant and utilizing bacteria. Glyphosate exposure induced only limited changes in the clam microbiome, with differences primarily detected in phylogenetic metrics and little evidence of broad shifts in diversity of community composition, suggesting that exposure effects were concentrated among less abundant members of the community. There was also little differentiation among the foot, gill, and digestive gland tissues sampled. *C. fluminea* was found to harbor bacteria capable of tolerating high concentrations of glyphosate, and eight cultured isolates demonstrated the ability to utilize glyphosate as a phosphorus source, with *phnJ* detection supporting involvement of the C-P lyase pathway.

Though these isolates represented minor members of the microbiome, further analysis of exposure and tissue-specific microbiomes revealed more abundant taxa belonging to groups with previously documented glyphosate-degrading capabilities. However, the low sample size and short exposure duration may not reflect the full effects of glyphosate on *C. fluminea* microbiome composition and diversity. Additionally, although our results demonstrate glyphosate utilization among cultured isolates, glyphosate reduction was not directly quantified and further investigation is needed to establish their degradation capacity.

Future studies should determine whether the limited microbiome response observed here is sustained across larger sample sizes, longer exposure periods, and populations of *C. fluminea* from habitats with different histories of herbicide exposure. Continued sampling could also help resolve the weak tissue differentiation observed in this study and determine whether it is characteristic of *C. fluminea* or reflects individual host variation. Further characterization of glyphosate-utilizing isolates should directly quantify glyphosate degradation and resulting metabolic products and determine whether these capabilities extend to other members of the clam microbiome. Together, these approaches would clarify the ecological significance of glyphosate-utilizing bacteria within freshwater bivalve microbiomes and determine whether bivalve-associated microbial communities may serve as sources of bacteria with potential applications in glyphosate bioremediation.

## Use of Generative AI

The authors used ChatGPT to assist with general grammar and spelling review. All manuscript content, scientific interpretation, and conclusions were developed and verified by the authors. OpenAI Codex was also used to assist with code execution and figure generation by facilitating workflows between QIIME2 and R command-line environments. All code, analyses, and figures were reviewed and verified by the authors.

## Data Availability

Raw sequencing data is publicly available on the National Center for Biotechnology Information (NCBI). 16S rRNA gene amplicon sequencing data is accessible through NCBI Sequence Read Archive, BioProject ID: PRJNA1541416. 16S rRNA sequences for all 10 isolates were deposited in the NCBI GenBank database under accession no. QB144937-QB144946.

## Author Contributions

SE conceived, designed, and executed the project; conducted analyses; and drafted the manuscript. CC conducted analyses. JPG executed the project. RED contributed to conception, project design, project execution, and manuscript drafting. All authors reviewed and approved the final manuscript.

## Acknowledgments

The authors would like to thank all members of the research team for their contributions to this work. Additional thanks are given to the Nonconnah Creek Conservancy in Memphis, TN for help with background and field site planning.

