## Supplemental Figures and Tables for "Glyphosate exposure minimally alters the *Corbicula fluminea* microbiome while associated bacteria exhibit high tolerance and degradation potential"

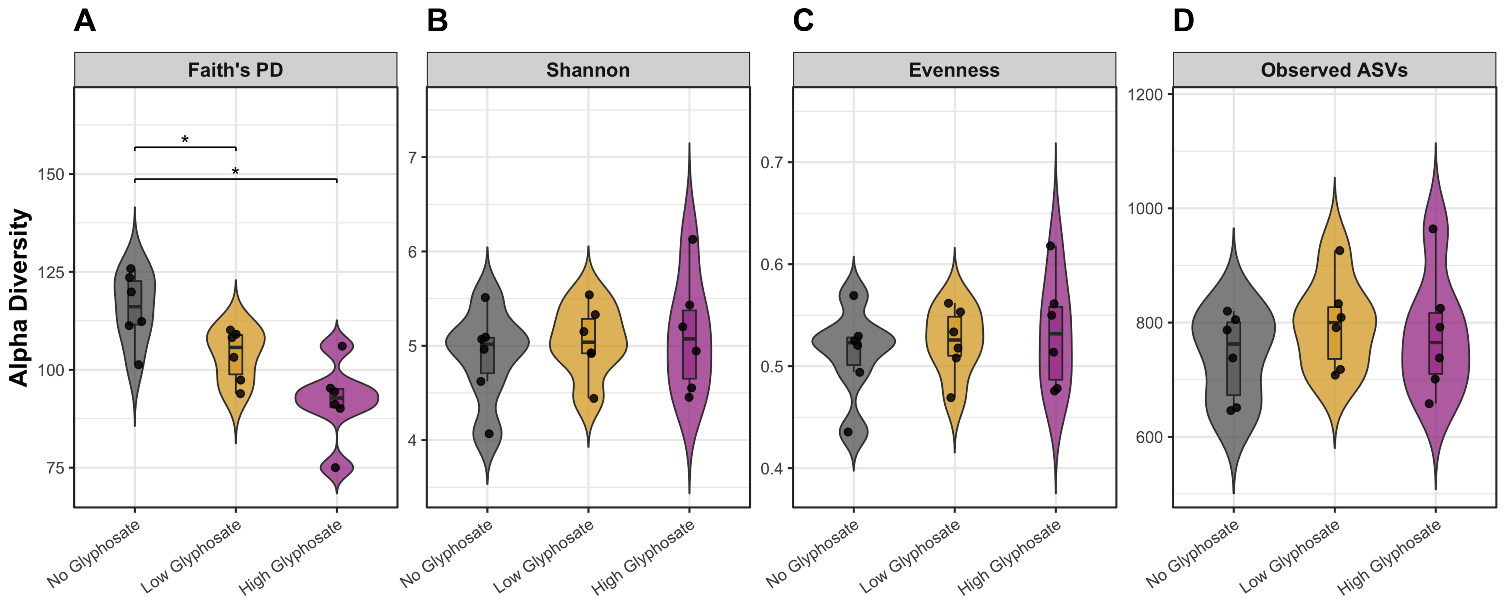


**Supplementary Figure 1**. Alpha diversity metrics associated with *Corbicula fluminea* microbial communities from the glyphosate exposure experiment. Diversity metrics examined were Faith’s PD, Shannon, Evenness, and observed exact sequence variants (ASVs). * denotes statistically significant differences between groups (q < 0.05).

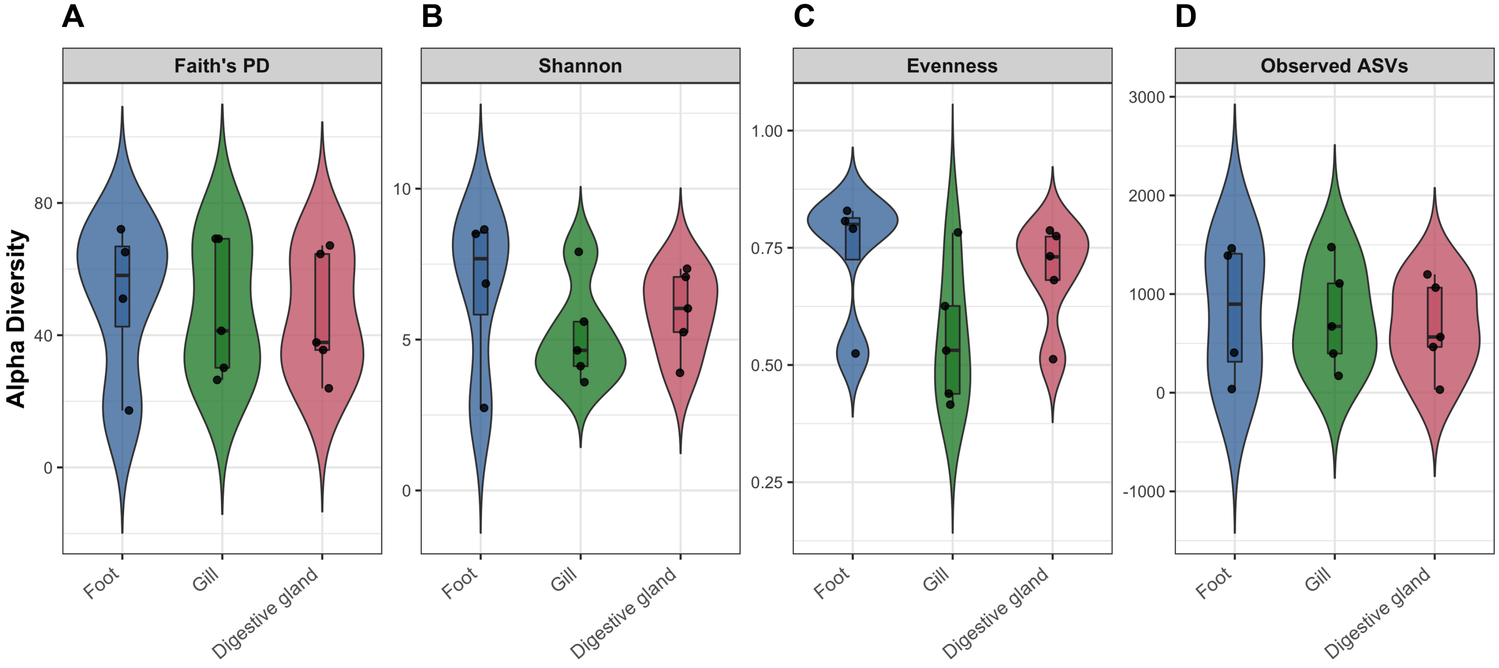


**Supplementary Figure 2.** Alpha diversity metrics associated with *Corbicula fluminea* microbial communities from the tissue-specific microbiomes from which bacteria were isolated. Diversity metrics examined were Faith's PD, Shannon, Evenness, and observed exact sequence variants (ASVs). * denotes statistically significant differences between groups ($q < 0.05$).

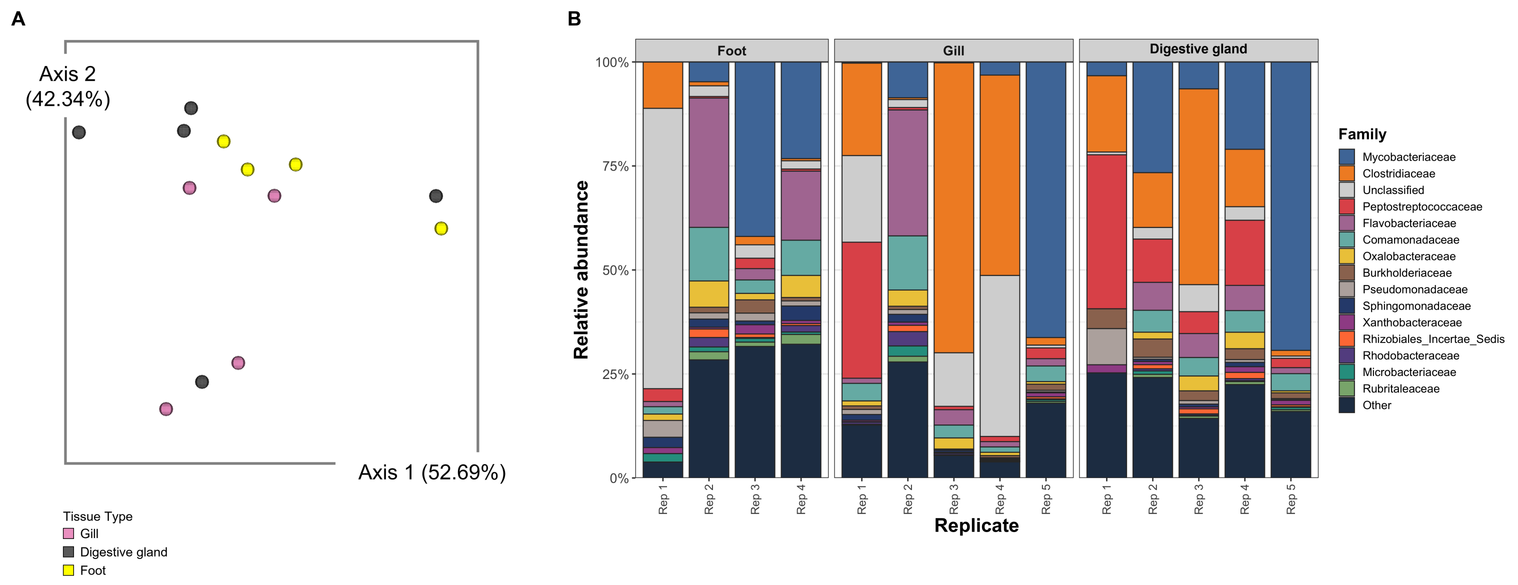


**Supplementary Figure 3.** A) Emperor biplot visualizing microbiome beta diversity from tissue-specific clam samples using robust Aitcheson Principal Component Analysis (RPCA). Individual gill, digestive gland, and foot tissues were used for microbiome analysis and bacteria isolation. B) Family level taxonomy bar plot for tissue-specific *Corbicula fluminea* microbiomes from which bacteria were isolated. Each bar represents an individual biological replicate. Samples are from individual gill, digestive gland, and foot tissue.

**Additional Supplementary Data Files:**

**Supplementary Table 1.** 16S rRNA gene sequences (1000 bp) and *phnJ* gene sequences (450 bp) for each of the 10 cultured isolates. (STable1_16S_phnj_Sequences.pdf).

| Isolate ID | Species identification | 16S rRNA sequence (5′–3′) | phnJ sequence (5′–3′) |
|---|---|---|---|
| Iso-42 | *Lelliottia amnigena* | NTGCAGTCGAGCGGTAGCACAGAAGAGCTTGCTCTC<br>GGGTGACGAGCGGCGGACGGGTGAGTAATGTCTGGG<br>AAACTGCCTGATGGAGGGGGATAACTACTGGAAACG<br>GTAGCTAATACCGCATAACGTCGCAAGACCAAAGAG<br>GGGGACCTTCGGGCCTCTTGCCATCAGATGTGCCCA<br>GATGGGATTAGCTAGTAGGTGGGGTAATGGCTCACC<br>TAGGCGACGATCCCTAGCTGGTCTGAGAGGATGACC<br>AGCCACACTGGAACTGAGACACGGTCCAGACTCCTA<br>CGGGAGGCAGCAGTGGGGAATATTGCACAATGGGCG<br>CAAGCCTGATGCAGCCATGCCGCGTGTATGAAGAAG<br>GCCTTCGGGTTGTAAAGTACTTTCAGCGAGGAGGAA<br>GGCATTGTGGTTAATAACCACAGTGATTGACGTTAC<br>TCGCAGAAGAAGCACCGGCTAACTCCGTGCCAGCAG<br>CCGCGGTAATACGGAGGGTGCAAGCGTTAATCGGAA<br>TTACTGGGCGTAAAGCGCACGCAGGCGGTCTGTCAA<br>GTCGGATGTGAAATCCCCGGGCTCAACCTGGGAACT<br>GCATTCGAAACTGGCAGGCTAGAGTCTTGTAGAGGG<br>GGGTAGAATTCCAGGTGTAGCGGTGAAATGCGTAGA<br>GATCTGGAGGAATACCGGTGGCGAAGGCGGCCCCCT<br>GGACAAAGACTGACGCTCAGGTGCGAAAGCGTGGGG<br>AGCAAACAGGATTAGATACCCTGGTAGTCCACGCCG<br>TAAACGATGTCGACTTGGAGGTTGTTCCCTTGAGGA<br>GTGGCTTCCGGAGCTAACGCGTTAAGTCGACCGCCT<br>GGGGAGTACGGCCGCAAGGTTAAAACTCAATGAATT<br>GACGGGGGCCCGCACAAGCGGTGGAGCATGTGGTTT<br>AATTCGATGCAACGCGAAGAACCTTACCTACTCTTG<br>ACATCCAGAGAACTTAGCAGAGATGCTTTGGTGCCT<br>TCGGGACTCTGAGACAGTGCTGCATGGC | NTTTTCCTNNNGNNNNATTGCCNGAACCNCTGCA<br>TTTTATCTATCCACGNNGAACGGAANCCCNCACC<br>ATGCACACTCTGGAATATTACGGTNATCATGCAC<br>GTGAAACTGGACNANGATATCNCCCGCTTCAGCC<br>ATATCTCCACCACCTACTCCTATCCNGAGAAGGT<br>GAACGGACACTACGTCATGGACCCGTCACCCATC<br>CCAAAATTCGATAACCGAAAATGGACATGATGCC<br>CGCGCTCCAGCTGTTTGGTGCCGGACGCGAAAAG<br>CGCATCTACGCCGTGCCGCCGTATACCCGCGTAG<br>AGAGTCTCGATTTTGACGATCATCCGTTCACGGT<br>GCAGGAGTGGGACGAGCCGTGCGCCATCTGCGGC<br>TCAACACACAGCTATCTCGACGAAGTGGTGCTGG<br>ACGATACGGGCAAACGGATGTTTGTCTGCTCCGA<br>ACCGAACT |
| Iso-50 | *Ewingella americana* | TNCAGTCGAGCGGCAGCGGAAGTANCNNGNTNCTTT<br>GCCGGCGAGCGGCGGACGGGTGAGTAATGTCTGGGA<br>AACTGCCTGATGGAGGGGGATAACTACTGGAAACGG<br>TAGCTAATACCGCATGACCTCGAAAGAGCAAAGTGG<br>GGGACCTTCGGGCCTCACGCCATCGGATGTGCCCAG<br>ATGGGATTAGCTAGTAGGTGAGGTAATGGCTCACCT<br>AGGCGACGATCCCTAGCTGGTCTGAGAGGATGACCA<br>GCCACACTGGAACTGAGACTGGTTCCACACTACTAC<br>GGGAGGCAGCAGTGGGGAATATTGCACAATGGGCGC<br>AAGCCTGATGCAGCCATGCCGCGTGTGTGAAGAAGG<br>CCTTCGGGTTGTAAAGCACTTTCAGCGAGGAGGAAG<br>GCGTTAAGGTTAATAACCTTGGCGATTGACGTTACT<br>CGCAGAAGAAGCACCGGCTAACTCCGTGCCAGCAGC<br>CGCGGTAATACGGAGGGTGCAAGCGTTAATCGGAAT<br>TACTGGGCGTAAAGCGCACGCAGGCGGTTTGTTAAG<br>TCAGATGTGAAATCCCCGAGCTTAACTTGGGAACTG<br>CATTTGAAACTGGCAAGCTAGAGTCTTGTAGAGGGG<br>GGTAGAATTCCAGGTGTAGCGGTGAAATGCGTAGAG<br>ATCTGGAGGAATACCGGTGGCGAAGGCGGCCCCCTG<br>GACAAAGACTGACGCTCAGGTGCGAAAGCGTGGGGA<br>GCAAACAGGATTAGATACCCTGGTAGTCCACGCTGT<br>AAACGATGTCGATTTGGAGGTTGTGGGCTTGACCCG<br>TGGCTTCCGGAGCTAACGCGTTAAATCGACCGCCTG<br>GGGAGTACGGCCGCAAGGTTAAAACTCAAATGAATT<br>GACGGGGGCCCGCACAAGCGGTGGAGCATGTGGTTT<br>AATTCGATGCAACGCGAAGAACCTTACCTACTCTTG<br>ACATCCAGAGAATTCGCTAGAGATAGCTTAGTGCCT<br>TCGGAACTCTGAGACAGTGCTGCATGGC | TNNTTTTCNAGNGCCCATCCCAGAGCCGCTGCGC<br>TTTATCGAGCCGCGCGAAACCGAAACGCGCAAGA<br>TGCACGCGCTGGAAGAGTACGGCGTGATGCAGGT<br>GAAATTGTATGAGGACATCGCGCGCTTCGGCCAT<br>ATCGCCACCACCTACGCCTATCCGGTCAAGGTGA<br>ACGATCGCTACGTGATGGACCCGTCGCCAATCCC<br>GAAATTTGATAACCCGAAAATGCACATGATGCCC<br>GCGCTGCAACTGTTCGGCGCTGGTCGCGAGAAAC<br>GCATTTACGCCCTGCCGCCGTTCACCAAGGTGGA<br>GAGCCTCGACTTTGAGGACCACCCGTTCACCGTG<br>CAGCAGTGGGACGAGCCTTGCGCTATTTGCGGCT<br>CGCACCACAGCTATCTCGACGAAGTGGTGCTCGA<br>CGACCTCGGCAACCGAATGTTCGTCTGCTCCGAT<br>ACCGGAAC |

| Isolate ID | Species identification | 16S rRNA sequence (5′–3′) | phnJ sequence (5′–3′) |
|---|---|---|---|
| Iso-88 | *Citrobacter freundii* | NTGCAGTCGANNGTAGCACNGNNGGAGCTTGCTCCT<br>TGGGTGACGAGTGGCGGACGGGNNNGTAATGTCTGG<br>GAAACTGCCCGATGGAGGGGGATAACTACTGGAAAC<br>GGTAGCTAATACCGCATAACGTCGCAAGACCAAAGA<br>GGGGGACCTTCGGGCCTCTTGCCATCGGATGTGCCC<br>AGATGGGATTAGCTAGTAGGTGGGGTAACGGCTCAC<br>CTAGGCGACGATCCCTAGCTGGTCTGAGAGGATGAC<br>CAGCCACACTGGAACTGAGACACGGTCCAGACTCCT<br>ACGGGAGGCAGCAGTGGGGAATATTGCACAATGGGC<br>GCAAGCCTGATGCAGCCATGCCGCGTGTATGAAGAA<br>GGCCTTCGGGTTGTAAAGTACTTTCAGCGAGGAGGA<br>AGGTGCTGTGGTTAATAACCGCAGCAATTGACGTTA<br>CTCGCAGAAGAAGCACCGGCTAACTCCGTGCCAGCA<br>GCCGCGGTAATACGGAGGGTGCAAGCGTTAATCGGA<br>ATTACTGGGCGTAAAGCGCACGCAGGCGGTCTGTCA<br>AGTCGGATGTGAAATCCCCGGGCTCAACCTGGGAAC<br>TGCATCCGAAACTGGCAGGCTAGAGTCTTGTAGAGG<br>GGGGTAGAATTCCAGGTGTAGCGGTGAAATGCGTAG<br>AGATCTGGAGGAATACCGGTGGCGAAGGCGGCCCCC<br>TGGACAAAGACTGACGCTCAGGTGCGAAAGCGTGGG<br>GAGCAAACAGGATTAGATACCCTGGTAGTCCACGCC<br>GTAAACGATGTCGACTTGGAGGTTGTGCCCTTGAGG<br>CGTGGCTTCCGGAGCTAACGCGTTAAGTCGACCGCC<br>TGGGGAGTACGGCCGCAAGGTTAAAACTCAAATGAA<br>TTGACGGGGGCCCGCACAAGCGGTGGAGCATGTGGT<br>TTAATTCGATGCAACGCGAAGAACCTTACCTACTCT<br>TGACATCCACAGAACTTGGCAGAGATGCCATGGTGC<br>CTTCGGGAACTNTGAGACAGTGCTGCAT | TCTTCCNGGTCCCGATCCCCGAGCCGTTGCGCTT<br>TATCGAGCCGCGCGAAACGGAAACCCGCACCATG<br>CACGCGCTGGAAGAGTACGGCATCATGCAGGTGA<br>AACTGTATGAAGATATCGCCCGCTTCGGCCATAT<br>CGCCACTACTTACGCCTATCCGGTAAAGGTCAAC<br>GACCGCTACGTCATGGACCCGTCGCCAATTCCAA<br>AATTCGACAATCCGAAAATGGATATGATGCCGGC<br>CCTGCAACTGTTTGGCGCAGGCCGTGAAAAGCGC<br>ATCTATGCCGTTCCACCTTACACCCGCGTGGAAA<br>GCCTCGATTTCGACGATCACCCGTTTACCGTGCA<br>GAGCTGGGATGAGCCTTGCGCCATCTGTGGCTCA<br>ACCCACAGCTATCTCGATGAAGTGGTGCTCGATG<br>ACACCGGCAAACGCATGTTTGTCTGCTCCGATAC<br>CGGANNNA |
| Iso-89 | *Atlantibacter hermannii* | NTGCAGTCGTANCANNNNNANCNCNNNCAGCTTGCT<br>GCTTTGCTGACGAGTGGCGGACGGGTGAGTAATGTC<br>TGGGAAACTGCCTGATGGCGGGGGATAACTACTGGA<br>AACGGTAGCTAATACCGCATAACGTCGCAAGACCAA<br>AGAGGGGGACCTTCGGGCCTCTTGCCATCGGATGTG<br>CCCAGATGGGATTAGCTAGTAGGTGGGGTAACGGCT<br>CACCTAGGCGACGATCCCTAGCTGGTCTGAGAGGAT<br>GACCAGCCACACTGGAACTGAGACACGGTCCAGACT<br>CCTACGGGAGGCAGCAGTGGGGAATATTGCACAATG<br>GGCGCAAGCCTGATGCAGCCATGCCGCGTGTATGAA<br>GAAGGCCTTCGGGTTGTAAAGTACTTTCAGCGGGGA<br>GGAAGGCGATACGGTTAATAACCGTGTCGATTGACG<br>TTACCCGCAGAAGAAGCACCGGCTAACTCCGTGCCA<br>GCAGCCGCGGTAATACGGAGGGTGCAAGCGTTAATC<br>GGAATTACTGGGCGTAAAGCGCACGCAGGCGGTCTG<br>TCAAGTCGGATGTGAAATCCCCGGGCTCAACCCGGG<br>AACTGCATTCGAAACTGGCAGGCTAGAGTCTTGTAG<br>AGGGGGGTAGAATTCCAGGTGTAGCGGTGAAATGCG<br>TAGAGATCTGGAGGAATACCGGTGGCGAAGGCGGCC<br>CCCTGGACAAAGACTGACGCTCAGGTGCGAAAGCGT<br>GGGGAGCAAACAGGATTAGATACCCTGGTAGTCCAC<br>GCCGTAAACGATGTCGACTTGGAGGTTGTGCCCTTG<br>AGGCGTGGCTTCCGGAGCTAACGCGTTAAGTCGACC<br>GCCTGGGGAGTACGGCCGCAAGGTTAAAACTCAAAT<br>GAATTGACGGGGGCCCGCACAAGCGGTGGAGCATGT<br>GGTTTAATTCGATGCAACGCGAAGAACCTTACCTGG<br>TCTTGACATCCACAGAATCCTGCAAAGATGCGGGAG<br>TGCCTTCGGAACTGTGAGACAGTGCTGC | ATTTTNCAGGNGCCNATCCCTGAAGCCGCTTGCG<br>CTTTATTGAACCGCGTGAAACGGAAACCCGCAAA<br>ATGNNCGCGCTGGAAAAGTACGGCGGGATGCAGG<br>TGAAACTGTATGAAAACATCGCGCGCTTCGGGCA<br>TATCGCCACCACTTACGCCTATCCGGGGAAAGTG<br>AATGACCGCTACGTGATGGATCCGTCGCCGATCC<br>CGAAATTTGATAACCCGAAAATGAACATGATGCC<br>CGCGTTACAGCTGTTCGGGGCCGGACGTGAAAAA<br>CGTATTTACGCCCTGCCGCCGTTCACCAAAGTCG<br>AAAGTCTGGATTTCAACAATCATCCTTTCACCGT<br>CCAGCAATGGGATGAACCCTGCGCGCTCTGCGGT<br>TCACACCACAGTTATCTCAATGAAGGGGTACTCA<br>ACAATCAGGGCAACCGCATGTTCGTCTGCTCCAA<br>ACCGNNNT |

| Isolate ID | Species identification | 16S rRNA sequence (5′–3′) | phnJ sequence (5′–3′) |
|---|---|---|---|
| Iso-96 | *Klebsiella pneumoniae* | NNTGCAGTCGAGCGGTAGCACAGAGAGCTTGCTCCT<br>CCGGGTGACGAGCGGCGGACGGGTGAGTAATGTCTG<br>GGAAACTGCCTGATGGAGGGGGATAACTACTGGAAA<br>CGGTAGCTAATACCGCATAACGTCGCAAGACCAAAG<br>TGGGGGACCTTCGGGCCTCATGCCATCAGATGTGCC<br>CAGATGGGATTAGCTAGTAGGTGGGGTAACGGCTCA<br>CCTAGGCGACGATCCCTAGCTGGTCTGAGAGGATGA<br>CCAGCCACACTGGAACTGAGACACGGTCCAGACTCC<br>TACGGGAGGCAGCAGTGGGGAATATTGCACAATGGG<br>CGCAAGCCTGATGCAGCCATGCCGCGTGTGTGAAGA<br>AGGCCTTCGGGTTGTAAAGCACTTTCAGCGGGGAGG<br>AAGGCGGTGAGGTTAATAACCTCATCGATTGACGTT<br>ACCCGCAGAAGAAGCACCGGCTAACTCCGTGCCAGC<br>AGCCGCGGTAATACGGAGGGTGCAAGCGTTAATCGG<br>AATTACTGGGCGTAAAGCGCACGCAGGCGGTCTGTC<br>AAGTCGGATGTGAAATCCCCGGGCTCAACCTGGGAA<br>CTGCATTCGAAACTGGCAGGCTAGAGTCTTGTAGAG<br>GGGGGTAGAATTCCAGGTGTAGCGGTGAAATGCGTA<br>GAGATCTGGAGGAATACCGGTGGCGAAGGCGGCCCC<br>CTGGACAAAGACTGACGCTCAGGTGCGAAAGCGTGG<br>GGAGCAAACAGGATTAGATACCCTGGTAGTCCACGC<br>CGTAAACGATGTCGATTTGGAGGTTGTGCCCTTGAG<br>GCGTGGCTTCCGGAGCTAACGCGTTAAATCGACCGC<br>CTGGGGAGTACGGCCGCAAGGTTAAAACTCAAATGA<br>ATTGACGGGGGCCCGCACAAGCGGTGGAGCATGTGG<br>TTTAATTCGATGCAACGCGAAGAACCTTACCTGGTC<br>TTGACATCCACAGAACTTTTCAGAGATGGATTGGTG<br>CCTTCGGGACTGTGAGACAGGTGCTGCA | NNNTCTACCAGGNGCCGATCCCGGAGCCATTGCG<br>CTTTATCGAGCCGCGTGAAACCGAAACCCGCACC<br>ATGCACGCCCTGGAAGAGTACGGCGTCATGCAGG<br>TGAAACTGTATGAAGATATCGCCCGCTTCGGCCA<br>TATCGCCACCACCTACGCCTATCCGGTGAAGGTC<br>AATGGCCGCTACGTGATGGACCCTTCGCCGATCC<br>CGAAATTCGATAACCCGAAAATGCACATGATGCC<br>GGCGCTGCAGCTGTTCGGCGCCGGGCGCGAAAAG<br>CGGATCTACGCCGTGCCGCCCTATACCCCGGTGG<br>AGAGTCTCGATTTCGACGACCATTCCTTTACCGT<br>GCAGGAGTGGGATGAGCCCTGCGCCATCTGCGGC<br>TCGCGCCACAGCTACCTCGACGAAGTGGTGCTGG<br>ATGACAGCGGCCAGCGGATGTTTGTCTGCTCCGA<br>CACCGGAC |
| Iso-97 | *Pantoea agglomerans* | TGCAGTCGNACGGTAGCACAGAGAGCTTGGTCTCTC<br>GTGACNAGAGGGGGACGGGNGAGTAATGTCTGGGGA<br>TCTNCCCCATATAGGGGGATATCTACTGTAAACGGT<br>GGGTAATACCGCATAACGTCGCAAGACCNNAGAGGG<br>GGACCTTCGGGCCTCTCGCCATCTCATGTGCCCCCA<br>TGGGATTATCTATAAGGTGGGGTAACGGCCCCCCTA<br>TGCGACAATCCCTATCTGTGCTGAGAGGATGACCAC<br>CCACACTGTAACTGAGACACGGTCCACACTCCTACA<br>GGAGGCAGCAGTGGGGAATATTGCGCAATGGGCGCA<br>AGCCTGATGCACCCATGCCGCGTGTGTGAAAAAAGC<br>CTTCTGGTTGTAAAGAACTTTCTCCGGGGAGGAAAG<br>CGATGTGGGTAATAACCCCGTCGATTGACNTTACCC<br>CCACAAAAAACACCGCNTATCTCCNNGCCACNANCC<br>GCGGNAATACNNAGGGGGCGAGCGTTAATCTNAATT<br>ACTGNGNGTAAAGCGCACGCACGCGGNCTGTGAANT<br>CACATGTGANATCNCCGNGNTCAACCTGTGAACTGT<br>GTTTGAGACTGTGANGNTNGAGTCTCGTANAGAGGG<br>GTANAATNCCANGTGTANCGNNGAAATGCGTANAGA<br>TCTGNAGGAATANNNNTGNGGAAAGCGGNCCCCTGT<br>ACNAAGANTGNGNCTCTNGTGCGANAGAGTGNGGAG<br>AAAACACGATNATATACNNTGTGANNCCACGCCGCA<br>NACNATGTCNANNTNNNANGNNGNNNCNCCTNNNAG<br>GNNNGNNNTNCNCNNAGNTANNNNNNNANAATCGNN<br>CNGCCNNNNGGAGNANNNNNNNCNANNNNNNAANNC<br>TNNNNNNNNNNNNGNNNGGGNCCNCNNCANNNGNNG<br>GNAGNATNTGNNTTATTNNTATGTNNNNNGANNNNN<br>TCATCNTNNNNNTCGTNNTCNNCANNANNNTTCNNA<br>NATGNNTTNNNGNCNNNNNNNNNNNNTN | TNNTCTACCAGGNGCCGATCCCGGAGCCATTGCG<br>CTTTATCGAGCCGCGTGAAACCGAAACCCGCACC<br>ATGCACGCCCTGGAAGAGTACGGCGTCATGCAGG<br>TGAAACTGTATGAAGATATCGCCCGCTTCGGCCA<br>TATCGCCACCACCTACGCCTATCCGGTGAAGGTC<br>AATGGCCGCTACGTGATGGACCCTTCGCCGATCC<br>CGAAATTCGATAACCCGAAAATGCACATGATGCC<br>GGCGCTGCAGCTGTTCGGCGCCGGGCGCGAAAAG<br>CGGATCTACGCCGTGCCGCCCTATACCCCGGTGG<br>AGAGTCTCGATTTCGACGACCATTCCTTTACCGT<br>GCAGGAGTGGGATGAGCCCTGCGCCATCTGCGGC<br>TCGCGCCACAGCTACCTCGACGAAGTGGTGCTGG<br>ATGACAGCGGCCAGCGGATGTTTGTCTGCTCCGA<br>TACCGANN |

| Isolate ID | Species identification | 16S rRNA sequence (5′–3′) | phnJ sequence (5′–3′) |
|---|---|---|---|
| Iso-98 | *Rahnella aquatilis* | NNTGCAGTCGAGCGGCAGCGGAAGTAGCTTGCTACT<br>TTGCCGGCGAGCGGCGGACGGGTGAGTAATGTCTGG<br>GAAACTGCCTGATGGAGGGGGATAACTACTGGAAAC<br>GGTAGCTAATACCGCATGACCTCGAAAGAGCAAAGT<br>GGGGGATCTTCGGACCTCACGCCATCGGATGTGCCC<br>AGATGGGATTAGCTAGTAGGTGAGGTAATGGCTCAC<br>CTAGGCGACGATCCCTAGCTGGTCTGAGAGGATGAC<br>CAGCCACACTGGAACTGAGACACGGTCCAGACTCCT<br>ACGGGAGGCAGCAGTGGGGAATATTGCACAATGGGC<br>GCAAGCCTGATGCAGCCATGCCGCGTGTGTGAAGAA<br>GGCCTTAGGGTTGTAAAGCACTTTCAGCGAGGAGGA<br>AGGCATCACACTTAATACGTGTGGTGATTGACGTTA<br>CTCGCAGAAGAAGCACCGGCTAACTCCGTGCCAGCA<br>GCCGCGGTAATACGGAGGGTGCAAGCGTTAATCGGA<br>ATTACTGGGCGTAAAGCGCACGCAGGCGGTTTGTTA<br>AGTCAGATGTGAAATCCCCGCGCTTAACGTGGGAAC<br>TGCATTTGAAACTGGCAAGCTAGAGTCTTGTAGAGG<br>GGGGTAGAATTCCAGGTGTAGCGGTGAAATGCGTAG<br>AGATCTGGAGGAATACCGGTGGCGAAGGCGGCCCCC<br>TGGACAAAGACTGACGCTCAGGTGCGAAAGCGTGGG<br>GAGCAAACAGGATTAGATACCCTGGTAGTCCACGCT<br>GTAAACGATGTCGACTTGGAGGTTGTGCCCTTGAGG<br>CGTGGCTTCCGGAGCTAACGCGTTAAGTCGACCGCC<br>TGGGGAGTACGGCCGCAAGGTTAAAACTCAAATGAA<br>TTGACGGGGGCCCGCACAAGCGGTGGAGCATGTGGT<br>TTAATTCGATGCAACGCGAAGAACCTTACCTACTCT<br>TGACATCCACGGAATTCGCCAGAGATGGCTTAGTGC<br>CTTCGGGAACCGTGAGACAGTGCTGCAT | TCTACCAGGNGCCGATCCCGGAGCCATTGCGCTT<br>TATCGAGCCGCGTGAAACCGAAACCCGCACCATG<br>CACGCCCTGGAAGAGTACGGCGTCATGCAGGTGA<br>AACTGTATGAAGATATCGCCCGCTTCGGCCATAT<br>CGCCACCACCTACGCCTATCCGGTGAAGGTCAAT<br>GGCCGCTACGTGATGGACCCTTCGCCGATCCCGA<br>AATTCGATAACCCGAAAATGCACATGATGCCGGC<br>GCTGCAGCTGTTCGGCGCCGGGCGCGAAAAGCGG<br>ATCTACGCCGTGCCGCCCTATACCCCGGTGGAGA<br>GTCTCGATTTCGACGACCATTCCTTTACCGTGCA<br>GGAGTGGGATGAGCCCTGCGCCATCTGCGGCTCG<br>CGCCACAGCTACCTCGACGAAGTGGTGCTGGATG<br>ACAGCGGCCAGCGGATGTTTGTCTGCTCCGATAC<br>CCGANNNA |
| Iso-99 | *Rahnella aquatilis* | NTGCAGTCGAGCGGCAGCGGAAGTAGCTTGCTACTT<br>TGCCGGCGAGCGGCGGACGGGTGAGTAATGTCTGGG<br>AAACTGCCTGATGGAGGGGGATAACTACTGGAAACG<br>GTAGCTAATACCGCATGACCTCGAAAGAGCAAAGTG<br>GGGGATCTTCGGACCTCACGCCATCGGATGTGCCCA<br>GATGGGATTAGCTAGTAGGTGAGGTAATGGCTCACC<br>TAGGCGACGATCCCTAGCTGGTCTGAGAGGATGACC<br>AGCCACACTGGAACTGAGACACGGTCCAGACTCCTA<br>CGGGAGGCAGCAGTGGGGAATATTGCACAATGGGCG<br>CAAGCCTGATGCAGCCATGCCGCGTGTGTGAAGAAG<br>GCCTTAGGGTTGTAAAGCACTTTCAGCGAGGAGGAA<br>GGCATCACACTTAATACGTGTGGTGATTGACGTTAC<br>TCGCAGAAGAAGCACCGGCTAACTCCGTGCCAGCAG<br>CCGCGGTAATACGGAGGGTGCAAGCGTTAATCGGAA<br>TTACTGGGCGTAAAGCGCACGCAGGCGGTTTGTTAA<br>GTCAGATGTGAAATCCCCGCGCTTAACGTGGGAACT<br>GCATTTGAAACTGGCAAGCTAGAGTCTTGTAGAGGG<br>GGGTAGAATTCCAGGTGTAGCGGTGAAATGCGTAGA<br>GATCTGGAGGAATACCGGTGGCGAAGGCGGCCCCCT<br>GGACAAAGACTGACGCTCAGGTGCGAAAGCGTGGGG<br>AGCAAACAGGATTAGATACCCTGGTAGTCCACGCTG<br>TAAACGATGTCGACTTGGAGGTTGTGCCCTTGAGGC<br>GTGGCTTCCGGAGCTAACGCGTTAAGTCGACCGCCT<br>GGGGAGTACGGCCGCAAGGTTAAAACTCAAATGAAT<br>TGACGGGGGCCCGCACAAGCGGTGGAGCATGTGGTT<br>TAATTCGATGCAACGCGAAGAACCTTACCTACTCTT<br>GACATCCACGGAATTCGCCAGAGATGGCTTAGTGCC<br>TTCGGGACCGTGAGACAGTGCTGCATGG | NNGNNGGNGNNNNANCCCTGNNCNNTGGCTCTTT<br>NTCNAACCCCGGTGGAAACCCAANNNCNNAAAAT<br>GGNCNCNCTGGAAGAATAACGGGGGGTGCANGGG<br>AAACTGGNTTAAAAACTCNCCCCCTTCCGGCATT<br>TTCNCCCCACTTTCCCCTATTCGGGGGAAGTGAA<br>TGACCGCTACCTGGTGGATCCCTCCCCCATCCCC<br>AAATTTGATTACCCCAAAATGAACATGATGCCCG<br>CATTGGANCTGGTTGGGGGCGGAANNGAAAAACG<br>CATTTACCCCCTGGCGCCGTTCACCAAAGTCGAA<br>AGTCTGGATTTCCACCAACATCCGTTCACCGTTC<br>AACAATGGGATGAACCTTGGGCGATTTGCGGGTC<br>CCACCCCCNCTATCTCCATGAAGTGGGGCTCCAC<br>GATTAAGGGAACCCCATGGTCCTCTGCTTCCACA<br>CCCAATAT |

| Isolate ID | Species identification | 16S rRNA sequence (5′–3′) | phnJ sequence (5′–3′) |
| --- | --- | --- | --- |
| Iso-100 | *Rahnella aquatilis* | TGCNAGTCGAGNGGCAGCGGNAAGTAGCTNGCTACT<br>NTGCCGGCAAGCGGCGGACGGGTGAGTAATGTCTGG<br>GAAACTGCCTGATGGAGGGGGATAACTACTGGAAAC<br>GGTAGCTAATACCGCATGACCTCGTAAGAGCAAAGT<br>GGGGGACCTTCGGGCCTCACGCCATCGGATGTGCCC<br>ACATGGGATTAGCTAGTAGGTGAGGTAATGGCTCAC<br>CTAGGCGACGATCCCTAGCTGGTCTGAGAGGATGAC<br>CAGCCACACTGGAACTGAGACNGGATCCACACTAAT<br>ACGGGAGGCAGCAGTGGGGAATATTGCACAATGGGC<br>GCAAGCCTGATGCAGCCATGCCGCGTGTGTGAAGAA<br>GGCCTTAGGGTTGTAAAGCACTTTCAGCGAGGAGGA<br>AGGGTTCAGTGTTAATAGCACTGAACATTGACGTTA<br>CTCGCAGAAGAAGCACCGGCTAACTCCGTGCCAGCA<br>GCCGCGGTAATACGGAGGGTGCAAGCGTTAATCGGA<br>ATTACTGGGCGTAAAGCGCACGCAGGCGGTTTGTTA<br>AGTCAGATGTGAAATCCCCGAGCTTAACTTGGGAAC<br>TGCATTTGAAACTGGCAAGCTAGAGTCTTGTAGAGG<br>GGGGTAGAATTCCAGGTGTAGCGGTGAAATGCGTAG<br>AGATCTGGAGGAATACCGGTGGCGAAGGCGGCCCCC<br>TGGACAAAGACTGACGCTCAGGTGCGAAAGCGTGGG<br>GAGCAAACAGGATTAGATACCCTGGTAGTCCACGCT<br>GTAAACGATGTCGACTTGGAGGTTGTGCCCTTGAGG<br>CGTGGCTTCCGGAGCTAACGCGTTAAGTCGACCGCC<br>TGGGGAGTACGGCCGCAAGGTTAAAACTCAAATGAA<br>TTGACGGGGGCCCGCACAAGCGGTGGAGCATGTGGT<br>TTAATTCGATGCAACGCGAAGAACCTTACCTACTCT<br>TGACATCCAGAGAATTCGCTAGAGATAGCTTAGTGC<br>CTTCGGGAACTCTGAGACNGGTGCTGCA | TTTTTCNGGNGCCGATCCCTGAGCCGCTGCNCTT<br>TATTGAACCGCGTGAAACGGAAACCCGCAAAATG<br>CACGCGCTGGAAAAGTACGGNGGGATGCAGGTGA<br>AACTGTATGAAAACATCGCGCGCTTCGGGCATAT<br>CGCCACCACTTACGCCTATCCGGGGAAAGTGAAT<br>GACCGCTACGNGATGGATCCGTCGCCAATCCCAA<br>AATTTGATAACCCGAAAATGAACATGATGCCCGC<br>GTTACAGCTGTTCGGGGCCGGACGTGAAAAACGT<br>ATTTACGCCCTGCCGCCGTTCACCAAAGTCAAAA<br>GTCTGGATTTCAACAATCATCCTTTCACCGTCCA<br>GCAATGGGATGAACCCTGCGCGCTCTGCGGTTCA<br>CACCACAGCTATCTCAATAAAGTGGTACTCAACA<br>ATCAGGGCAACCGCATGTTCNTCTGCTCCAANNN<br>NNAATTAT |
| Iso-111 | *Lelliottia amnigena* | NTGCAGTCGAGCGGTAGCACAGAGAGCTTGCTTCTN<br>CGGGTGACGAGCGGCGGACGGGTGAGTAATGTCTGG<br>GAAACTGCCTGATGGAGGGGGATAACTACTGGAAAC<br>GGTAGCTAATACCGCATAACGTCGCAAGACCAAAGA<br>GGGGGACCTTCGGGCCTCTTGCCATCAGATGTGCCC<br>AGATGGGATTAGCTAGTAGGTGGGGTAATGGCTCAC<br>CTAGGCGACGATCCCTAGCTGGTCTGAGAGGATGAC<br>CAGCCACACTGGAACTGAGACACGGTCCAGACTCCT<br>ACGGGAGGCAGCAGTGGGGAATATTGCACAATGGGC<br>GCAAGCCTGATGCAGCCATGCCGCGTGTATGAAGAA<br>GGCCTTCGGGTTGTAAAGTACTTTCAGCGAGGAGGA<br>AGGCATTGTGGTTAATAACCACAGTGATTGACGTTA<br>CTCGCAGAAGAAGCACCGGCTAACTCCGTGCCAGCA<br>GCCGCGGTAATACGGAGGGTGCAAGCGTTAATCGGA<br>ATTACTGGGCGTAAAGCGCACGCAGGCGGTCTGTCA<br>AGTCGGATGTGAAATCCCCGGGCTCAACCTGGGAAC<br>TGCATTCGAAACTGGCAGGCTAGAGTCTTGTAGAGG<br>GGGGTAGAATTCCAGGTGTAGCGGTGAAATGCGTAG<br>AGATCTGGAGGAATACCGGTGGCGAAGGCGGCCCCC<br>TGGACAAAGACTGACGCTCAGGTGCGAAAGCGTGGG<br>GAGCAAACAGGATTAGATACCCTGGTAGTCCACGCC<br>GTAAACGATGTCGACTTGGAGGTTGTTCCCTTGAGG<br>AGTGGCTTCCGGAGCTAACGCGTTAAGTCGACCGCC<br>TGGGGAGTACGGCCGCAAGGTTAAAACTCAAATGAA<br>TTGACGGGGGCCCGCACAAGCGGTGGAGCATGTGGT<br>TTAATTCGATGCAACGCGAAGAACCTTACCTACTCT<br>TGACATCCAGAGAACTTAGCAGAGATGCTTTGGTGC<br>TTCGGGACTCTGAGACAGTGCTGCATGG | NNATTTTCCTNGNGNNNNATTGCCNGCANCNNCT<br>GCATTTTATCTAGCCACGCGNNAACGGAAACCCG<br>CACCATGCACACTCTGGAAGAGTACGGTGATGAT<br>GCACGTGAAACTGGACNANGATATCGCCCGCTTC<br>NGCCATATCTCCACCACCTACTCCTATCCCGAGA<br>AGGTGAACGGACACTACGTCATGGACCCGTCACC<br>CATCCCAAAATTCGATAACCCGAAAATGGACATG<br>ATGCCCGCGCTCCAGCTGTTTGGTGCCGGACGCG<br>AAAAGCGCATCTACGCCGTGCCGCCGTATACCCG<br>CGTAGAGAGTCTCGATTTTGACGATCATCCGTTC<br>ACGGTGCAGGAGTGGGACGAGCCGTGCGCCATCT<br>GCGGCTCAACACACAGCTATCTCGACGAAGTGGT<br>GCTGGACGATACGGGCAAACGGATGTTTGTCTGC<br>TCCGAACC |

**Supplementary Table 2.** Relative abundance of amplicon sequence variants (ASVs) matching bacterial isolate sequences recovered from *Corbicula fluminea* gill, digestive gland, and foot tissue. ASVs were identified by comparison with 16S rRNA gene sequences from cultured isolates. Values represent relative abundances (%) within the glyphosate exposure microbiome and tissue microbiome datasets. Matches represent the closest non-exact sequence matches to cultured isolates.

| | | Glyphosate Exposure Microbiome | | | Tissue Microbiome | | |
|---|---|---|---|---|---|---|---|
| **Isolate ID** | **Species name** | **No glyphosate relative abundance (%)** | **Low glyphosate relative abundance (%)** | **High glyphosate relative abundance (%)** | **foot** | **gill** | **dg** |
| Iso-42 | *Lelliotia amnigena* | 0.01139532 | 0.00796921 | 0.02645623 | 0.00000000 | 0.00000000 | 0.00228194 |
| Iso-50 | *Ewingella americana* | 0.00000000 | 0.00000000 | 0.00134915 | 0.00000000 | 0.00000000 | 0.00285242 |
| Iso-88 | *Citrobacter freundii* | 0.03275370 | 0.01799128 | 0.04512593 | 0.00000000 | 0.00000000 | 0.00570485 |
| Iso-89 | *Atlantibacter hermannii* | 0.00041460 | 0.00155694 | 0.00000000 | 0.00000000 | 0.00118177 | 0.00000000 |
| Iso-96 | *Klebsiella pneumoniae* | 0.00000000 | 0.00000000 | 0.00043687 | 0.00000000 | 0.00118177 | 0.00000000 |
| Iso-97 | *Pantoea agglomerans* | 0.00000000 | 0.00000000 | 0.00043687 | 0.00000000 | 0.00137873 | 0.00000000 |
| Iso-98 | *Rahnella aquatilis* | 0.00071613 | 0.00000000 | 0.00000000 | 0.00000000 | 0.00078784 | 0.0019967 |
| Iso-99 | *Rahnella aquatilis* | 0.00071613 | 0.00000000 | 0.00000000 | 0.00000000 | 0.00078784 | 0.0019967 |
| Iso-100 | *Rahnella aquatilis* | 0.00000000 | 0.00000000 | 0.00038547 | 0.02360917 | 0.00000000 | 0.00000000 |
| Iso-111 | *Lelliottia amnigena* | 0.01139532 | 0.00796921 | 0.02645623 | 0.00000000 | 0.00000000 | 0.00228194 |
